# Ultra-fast fit-free analysis of complex fluorescence lifetime imaging via deep learning

**DOI:** 10.1101/523928

**Authors:** Jason T. Smith, Ruoyang Yao, Nattawut Sinsuebphon, Alena Rudkouskaya, Joseph Mazurkiewicz, Margarida Barroso, Pingkun Yan, Xavier Intes

## Abstract

Fluorescence lifetime imaging (FLI) provides unique quantitative information in biomedical and molecular biology studies, but relies on complex data fitting techniques to derive the quantities of interest. Herein, we propose a novel fit-free approach in FLI image formation that is based on Deep Learning (DL) to quantify complex fluorescence decays simultaneously over a whole image and at ultra-fast speeds. Our deep neural network (DNN), named FLI-Net, is designed and model-based trained to provide all lifetime-based parameters that are typically employed in the field. We demonstrate the accuracy and generalizability of FLI-Net by performing quantitative microscopic and preclinical experimental lifetime-based studies across the visible and NIR spectra, as well as across the two main data acquisition technologies. Our results demonstrate that FLI-Net is well suited to quantify complex fluorescence lifetimes, accurately, in real time in cells and intact animals without any parameter settings. Hence, it paves the way to reproducible and quantitative lifetime studies at unprecedented speeds, for improved dissemination and impact of FLI in many important biomedical applications, especially in clinical settings.

## I. INTRODUCTION

MOLECULAR imaging has become an indispensable tool in biomedical studies with great impact on numerous fields from fundamental biological investigations to transforming clinical practice. Among all molecular imaging modalities, fluorescence optical imaging is a central technique thanks to its high sensitivity, the numerous molecular probes available, either endogenous or exogenous, and its ability to simultaneously image multiple biomarkers or biological processes at various spatio-temporal scales^1,2^. Especially, fluorescence lifetime imaging (FLI) has become an ever increasingly popular method as it provides unique insights into the cellular micro-environment, by non-invasively examining numerous intracellular parameters^3^ such as metabolic status^4^, reactive oxygen species^5^ and intracellular pH^6^. Moreover, FLI’s exploitation of native fluorescent signatures has been extensively investigated for enhanced diagnostic of numerous pathologies.^7–10^ FLI is also the most accurate approach to quantify Förster Resonance Energy Transfer (FRET), an invaluable technique used to quantify protein-protein interactions, biosensor activity and ligand-receptor engagement in vivo.^11^ FLI is not a direct imaging modality and beyond dedicated imaging platforms, the acquired temporal data set needs to be post-processed to quantify the lifetime or lifetime-based parameters. Such post-processing typically involves a model-based process in which iterative optimization methods are employed to estimate the different parameters of interest (mean-lifetime, FRET efficiencies or population fractions). Mono- or bi-exponential models, depending on the application at hand, are the most widely employed to analyze FLI datasets. Yet, it is notorious that the accuracy of these methods is often associated with user defined parameter settings employed to constrain the inverse problem. These methods are also relatively slow and/or computationally expensive.^12^ This complexity together with a lack of standardized methods has limited the widespread use and impact of FLI, especially clinically. Recently, a fit-free lifetime quantification methodology has been proposed, the phasor approach.^13^ The phasor method is a graphical representation of excited-state fluorescence lifetimes for *in vitro* systems. The phasor technique has been widely adopted due to its simplicity that allows non-imaging experts to perform simple and fit-free analyses of the information contained in the many thousands of pixels constituting an image^14^. However, although the phasor method provides a graphical interface that simplifies FLI data interpretation, the mathematics underlying its computation can be challenging. The approach needs to be modified for techniques such as time-gated fluorescence^15^ and typically requires some calibration samples to be quantitative^16^.

In parallel, interests in data driven and model-free processing of imaging methodologies has boomed over the last decade. Of particular note, Machine Learning (ML) and Deep Learning (DL) methods have recently profoundly impacted the image processing field. For example, deep-neural networks (DNNs) are currently providing high level, robust performances in numerous biomedical applications – such as in pathology through multiple imaging modalities,^17–19^ natural language processing,^20^ image reconstruction via direct mapping from the sensor-image domain^21^ and reinforcement learning applied to drug discovery.^22^ DL methods are increasingly employed in molecular optical imaging applications from resolution enhancement in histopathology^23^, super resolution microscopy^24^, fluorescence signal prediction from label-free images^25^, single molecular localization^26^, fluorescence microscopy image restoration^27^ and hyperspectral single pixel lifetime imaging for instance^28^. However, the typical application of DL methods to image processing is data driven and hence, requires large data sets that are either difficult to acquire and/or not readily available. Moreover, the performances of DL methods can be limited to the specific data set employed for training and hence, not generalized.

Herein, we present a 3D Convolutional Neural Network (CNN) that is designed to process the classical data sets acquired by current fluorescence imaging systems to provide, in quasi real-time, the lifetime maps as well as associated quantities (i.e.: mean-lifetime, fractional amplitude of fluorescence species, FRET Efficiency (E%) or FRET Donor Fraction (FD%)). Our 3D CNN approach, unlike previous methods, does not require any user-defined parameter entry, is able to tackle either mono- or bi-exponential data sets, is accurate for a large range of lifetimes (even ones close to the instrument response) and provides superior performances in photon starved conditions. Furthermore, the 3D CNN can be trained efficiently using a synthetic data generator and validated with experimental data sets, avoiding the need to acquire massive training datasets experimentally. Additionally, we demonstrate that our 3D CNN is capable of processing experimental fluorescent decays acquired by either Time Correlated Single Photon Counting (TCSCP)- or gated ICCD- based instruments, which are the two main technologies employed in the field. Herein, the potential of the proposed approach is demonstrated by performing FLI microscopy to quantify the metabolic status of live cells as well as reporting FRET to measure levels of receptor engagement. Moreover, the experimental demonstration includes applications in the visible as well as the near-infrared (NIR) range, allowing for a large range of lifetimes to be considered. In all cases, the 3D CNN performances are benchmarked against the widely used FLIM processing software *SPCImage*^29^. Lastly, we demonstrate the potential of our 3D CNN to quantify whole-body dynamic lifetime-based FRET occurrence in a live animal at unprecedented time frames (≌ 32ms per full whole-body image). Overall, these results demonstrate that DL methodologies, beyond classical image processing tasks, are well suited for image formation paradigms that to date were based on inverse problem solvers. Our reported 3D CNN architecture and training strategy provide a versatile and generalized new tool for fit-free analysis of complex fluorescence lifetime imaging processes. Due to its ease of use and ultra-fast qualities, our 3D CNN should further stimulate the widespread use of FLI techniques, provide standardized quantification capabilities (as no parameter settings are required) and enable new applications such as real-time wide-field FLI in pre-clinical and clinical studies, especially facilitating optical guided surgery.

## II. 3D-CNN ARCHITECTURE, TRAINING AND VALIDATION

**Figure 1.**
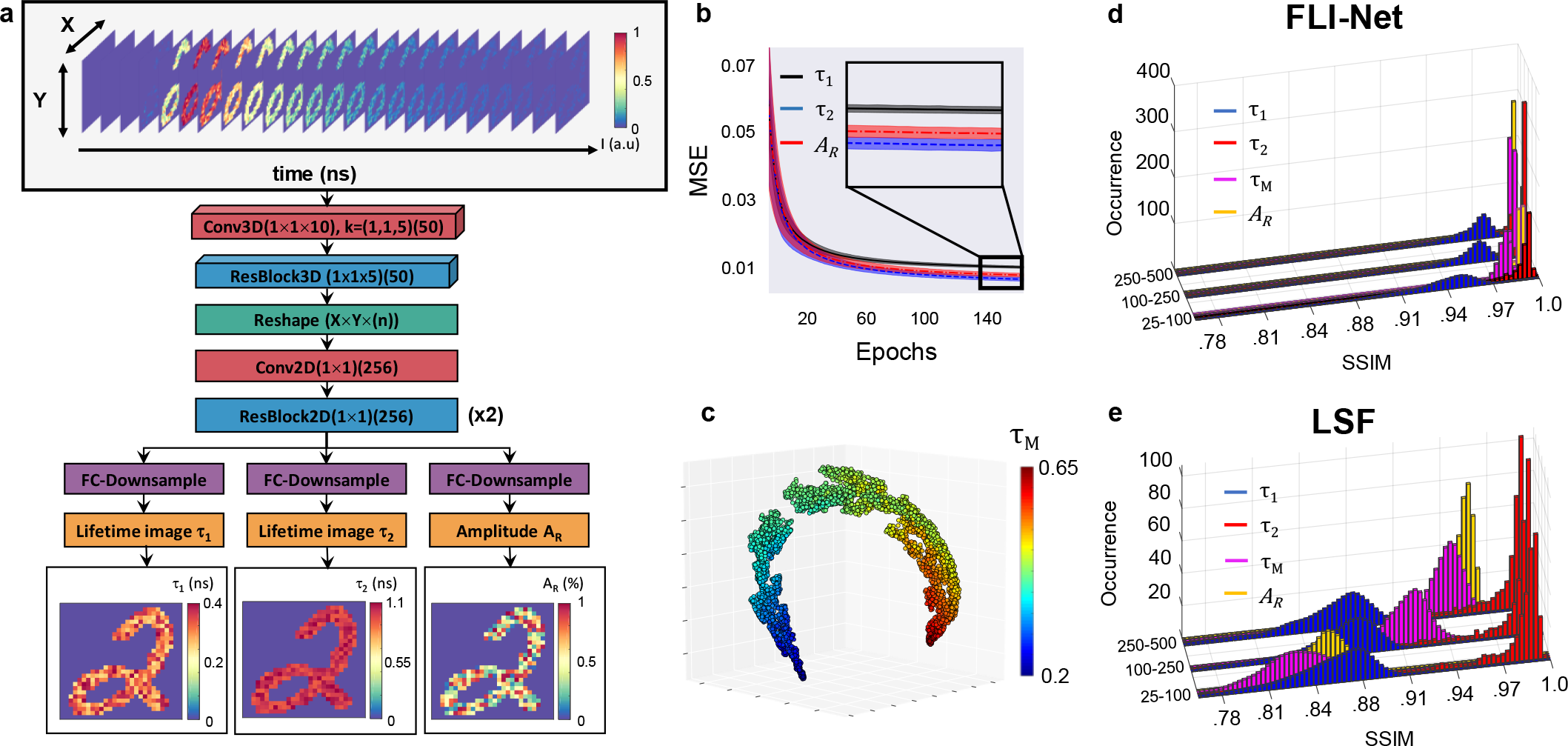
Illustration of our 3D-CNN (“FLI-Net”) structure and corresponding metrics of note. During the training phase, the input to our DNN (**a**) was a set of simulated data voxels containing a TPSF at every non-zero value of a randomly chosen MNIST image. After a series of spatially-independent 3D-convolutions, the voxel underwent a reshape (from 4D to 3D) and subsequently branched into three separate series of fully-convolutional downsampling for simultaneous three-image reconstruction. **b**, 30 MSE validation curve average with corresponding standard deviation (shaded) for each parameter. **c**, t-SNE visualization obtained via the last activation map before the tri-junction reconstruction, where each point represents a TPSF voxel assigned a randomized trio of lifetime and amplitude ratio values. FLI-Net performance (**d**) versus LSF (**e**) upon evaluation of simulated TPSF voxels over three ranges of maximum photon count.

The 3D CNN, named “FLI-Net” (Fluorescence Lifetime Imaging-Network) hereafter is designed to mimic a curve-fitting approach using layers of convolutional operations and non-linear activation functions. FLI-Net is designed such that time-and spatially resolved fluorescence decays are input as 3D data cube (*x*,*y*,*t*) and bi-exponential parameters (two lifetimes and one fractional amplitude) are estimated at each pixel to be provided in output images of the same dimension as the input (*x*,*y*). A rendering of the architecture of FLI-Net is provided in **Fig. 1a**.

The network architecture consists of two main parts: 1) a shared branch for temporal feature extraction and 2) a subsequent three-junction split into separate branches for simultaneous reconstruction of short lifetime (***τ***_**1**_), long lifetime (***τ***_**2**_) and fractional amplitude of the short lifetime (***A***_***R***_). There are a couple of design choices that are critical to the performance of FLI-Net; providing the basis for a consequently high level of sensitivity, stability, speed and reconstructive accuracy. First, it is crucial to introduce convolutions (Conv3D) along the temporal dimension at each spatially located pixel at the first layer in order to maximize spatially-independent feature extraction along each TPSF. After this step, a residual block (ResBlock) of reduced kernel length is employed. This second step enables further extraction of temporal information while reaping the benefits obtained through residual learning (elimination of vanishing gradients, no overall increase in computational complexity or parameter count, etc.)^30^. The beneficial implementation of residual learning has been thoroughly documented in not only image classification and segmentation^31^ but also in areas of speech recognition^32^. Fully-convolutional networks^33^, or networks designed such that input of any spatial dimensionality can be analyzed with no loss in performance, offer enormous benefit to problems where 1) prior knowledge of input size is inherently variable and 2) the experimental data of interest is memory exhaustive. After performing the common features of the whole input, the network splits into three dedicated branches to estimate the individual lifetime-based parameters of interest. In each of these branches, a sequence of convolutions is employed for down-sampling to the intended 2D image. A more detailed description of the network architecture is provided in the ***Methods*** section.

To obtain large data sets to train FLI-Net and validate its architecture robustness, feature extraction efficiency as well as quantitative accuracy, we generated 10,000 temporal point-spread function (TPSF) voxels using a bi-exponential model convolved with an experimental instrument response function (IRF). The parameters of the bi-exponential model were varied over the lifetime range considered in the application (Visible (***τ***_**1**_, ***τ***_**2**_)∈[0.2-3]ns; NIR (***τ***_**1**_, ***τ***_**2**_)∈[0.2-1.1]ns) and the fractional amplitude ***A***_***R***_ varied from 0 to 100% (***A***_***R***_ = 0 or 100% are corresponding to mono-exponential decays whereas every value between these extreme corresponds to bi-exponential decays). See Table S1 in the Supplementary Material section for a full summary of the parameters used for training. The IRF was acquired from our gated-ICCD. Last, the photon counts (p.c.) of the maximum of the TPSF were set between 250 and 2,000 counts followed by the addition of Poisson noise. The training data set was split into training (8,000) and validation (2,000) data sets. Additional information on the generation of this data set can be found in the ***Methods***. To demonstrate the robustness of FLI-Net, training and validation were performed over 30 times with randomly initialized training/validation partitions. The plotted average of 30 validation mean-squared error (MSE) curves trained over 150 epochs with corresponding standard deviation bounds for all three output branches is provided in **Fig. 1b** and illustrates the DNN’s excellent convergence stability. To evaluate if the feature extraction of the shared branch was robust and effective, we registered the output of the shared branch’s final activation layer during feed-through of 5,000 newly simulated TPSF data voxels (not used in training or validation). These high-dimension features were flattened and projected to a 3D feature space via t-SNE^34^. Their display as a scatter plot is provided in **Fig. 1c**. The continuous gradient observed in the 3D plot of the t-SNE values versus the mean lifetimes simulated ***τ***_***m***_ = ***A***_***R***_***τ***_**1**_ + (**1** − ***A***_***R***_)***τ***_**2**_; ***τ***_***m***_ ∈ ([0.2, 0.65] ns) indicates an efficient and sensitive feature extraction for lifetime-based parameter estimation. Beyond feature extraction, we provide also the summary of the quantitative accuracy of the network in estimating the three above mentioned lifetime-based parameters (***τ***_**1**_, ***τ***_**2**_, ***A***_***R***_). The accuracy of the results is evaluated via the Structural Similarity Index (SSIM) between the simulated and estimated values (SSIM=1 indicating perfect one-to-one concordance). The SSIMs are also reported for three ranges of maximum photon counts (i.e., p.c._good_∈[250-500]; p.c._challenging_∈[100-250]; p.c._low_∈[25,100]) as lifetime-based biomedical imaging is notoriously a photon-starved application. In all three cases, FLI-Net (**Fig. 1d**) significantly outperforms the classical LSF (**Fig. 1e**) method, which as expected, demonstrates worsening performances at very low photon counts 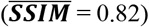. Note, that although the network was only trained using TPSF data possessing intensity values greater than 500 maximum photon counts in this specific instance, it performs extremely well for low photon counts levels too (even in the 25-100 range with a 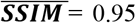 for the worse case). Overall, these training and validating results establish that FLI-Net can be efficiently and robustly trained via synthetic data representing both mono- and bi-exponential decays. Moreover, FLI-Net outperforms the classical LSF approach in estimating the three lifetime-based parameters that are commonly employed in FLI applications. To further establish the usefulness and unique potential of FLI-Net, we evaluated its performance when using experimental data sets after training with simulation data generated through the workflow previously described.

**Figure 2.**
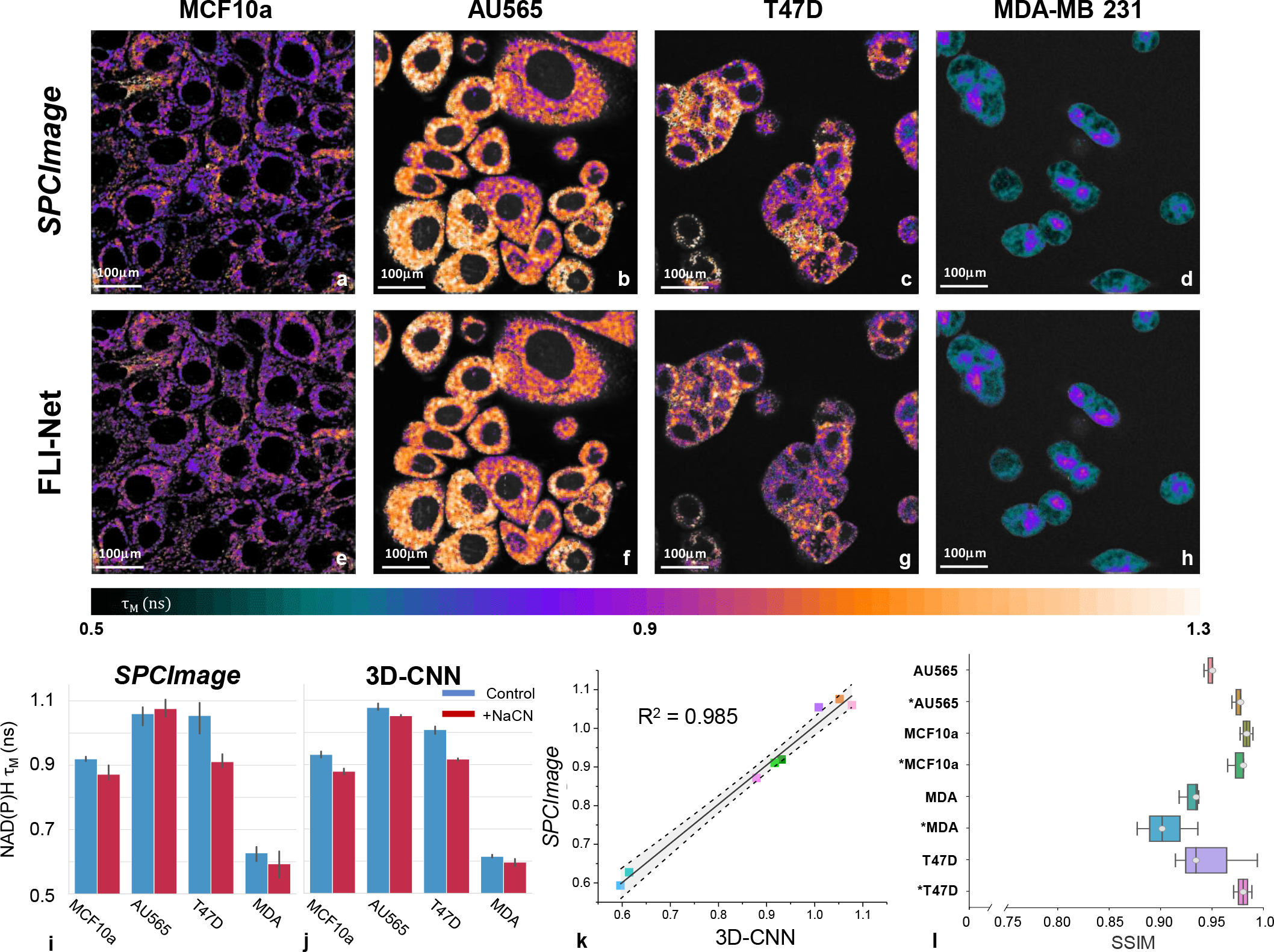
Visible FLIM microscopy data of NAD(P)H. **a-d**, Representative maps of NAD(P)H ***τ***_***m***_ obtained with the commercial software *SPCImage* and FLI-Net (e-h). **i-j**, Averaged NAD(P)H ***τ***_***m***_values obtained across all FLIM data using both techniques. **k**, Linear regression with corresponding 95% confidence band (grey shading) of averaged NAD(P)H ***τ***_***m***_ values from all cell-lines pre and post-exposure to Na cyanide (n = 3) obtained via *SPCImage* and our DNN (slope = 1.01 (SE = .05); p < 1e-5; intercept = −6.9e-3 (SE = 4.2e-2); R^2^ = 0.985). The microscopy data acquired post-exposure are notated with an asterisk. **l**, SSIM measurements for all NAD(P)H ***τ***_***m***_ images. Further metrics of note are included in the *Supplementary Material*.

## III. FLUORESCENCE LIFETIME IMAGING MICROSCOPY

Fluorescence lifetime imaging microscopy (FLIM) is the most widely used FLI application. For this study, we have selected metabolic and FRET imaging as they are some of the most challenging, yet sought after, FLIM applications. We report first the performance of FLI-Net in quantifying the metabolic status of live cells as reported by NAD(P)H imaging. Secondly, we report on FLI-Net’s accuracy in quantifying ligand-receptor engagement via lifetime-based FRET in the visible and NIR range.

### Metabolic imaging

Quantification of the fractional concentration between free and protein-bound NADH provides important information regarding cellular metabolic state. Given that both free and protein-bound NADH possess the same absorption and emission profiles, but differ significantly in fluorescence lifetime, FLIM has been used extensively for sensitive free vs. bound NADH quantification in vitro.^35^ First, confocal FLIM data were collected from four human cell lines (MCF10A as a non-cancerous mammary epithelial cell line, the remaining being cancer cell lines representing different types of breast cancer) using a Zeiss LSM 880 Airyscan NLO multiphoton confocal microscope equipped with HPM-100-40 high speed hybrid FLIM detector (GaAs 300-730 nm; Becker & Hickl) and a Titanium: Sapphire laser (Ti: Sa) (680-1040 nm; Chameleon Ultra II, Coherent, Inc.). These different cell lines have been shown to exhibit markedly different metabolic states, as reported by NADH ***τ***_***m***_.^36^ Additionally, the cells were exposed to 2.5 mM of Na cyanide (NaCN), which is a well-known inhibitor of many metabolic processes leading to reduced NADH ***τ***_***m***_^36^. FLIM acquisition was performed prior to exposure and after 30-minute incubation of live cells with NaCN. The FLIM NADH ***τ***_***m***_ images for each case are provided in **Fig. 2**, both for FLI-Net and *SPCImage*. A visual inspection of these images shows that FLI-Net and *SPCImage* provides strikingly similar results. The descriptive statistics of NADH ***τ***_***m***_ of *SPCImage* versus FLI-Net for each case is summarized in **Fig. 2i-l**. The excellent concordance between the two analytic frameworks is highlighted in **Fig. 2k** by very high coefficients of determination (R^2^∈[0.985]) and low p-value (p < 1e-5). Additionally, we provide the SSIM between FLI-Net and *SPCImage* in **Fig. 2l**. The SSIM values indicate an excellent spatial congruence between FLI-Net and *SPCImage* in all cases, with the lowest measured value obtained for MDA-NaCN 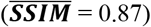.

### Ligand-receptor engagement

FLIM-FRET imaging enables the quantification of biosensor activity that can report on many cellular processes and/or ligand-receptor engagement and subsequent cellular internalization. FLIM-FRET is typically performed using visible light and by quantifying the reduction in the donor ***τ***_***m***_ associated with FRET quenching. Herein, visible FRET-FLI microscopy data were collected using a Zeiss LSM 510 equipped with HPM-100-40 high speed hybrid FLIM detector (GaAS 300-730 nm; Becker & Hickl) and a Titanium: Sapphire laser (Ti: Sa) (Chameleon) (apparatus, fluorescence labeling, and data acquisition details described elsewhere^37^). T47D human breast cancer cells were incubated with Tf-AF488 and Tf-AF555 visible FRET pair with a range of acceptor-donor (A:D) ratio from 0:1 to 2:1. As the A:D ratio increases, it is expected that FRET occurrence increases with the donor ***τ***_***m***_ decreasing accordingly. The ***τ***_***m***_ for each condition as estimated via FLI-Net are provided in **Fig. 3** (upper panel). In all cases, the ***τ***_***m***_ estimated are within the expected range and, overall, follow the decreasing trend expected as A:D ratio increases. To assess if FLI-Net provides similar results as *SPCImage*, we show in **Fig. 3c** and **Fig. 3d** the respective distribution of estimated ***τ***_***m***_. For the four A:D ratios reported, the mean-lifetime distributions are in excellent agreement between FLI-Net and *SPCImage*. Furthermore, we computed the Bhattacharyya coefficient (BC) to measure the similarity of these paired probability distributions. As displayed in **Fig. 3e** (further discussed in ***Methods***), the BCs are all very close to ≅ 1 indicating that FLI-Net and *SPCImage* provide almost identical ***τ***_***m***_ distributions for all cases. Additionally, we computed the MSE to assess spatial congruency between the two post-processing methods. As reported in **Fig. 3f**, the MSEs values are low, indicating a very good pixel-pixel correspondence.

Beyond the quantitative and spatial accuracy of FLI-Net as demonstrated by its benchmarking against *SPCImage*, we compared its computational speed versus *SPCImage* on the same computational platform. The time required for analysis of each TCSPC voxel, which possessed 256 time-points (visible FLIM) with a pixel resolution of 512 × 512, was just 2.5 seconds on average using our 3D CNN compared to ≅ 45 seconds with *SPCImage*. Though it is important to note that one input parameter of importance to *SPCImage* is the photon count restrictions that leads to fitting only a small subset of the pixel in the input voxel, whereas FLI-Net processes the voxel’s entirety. Taking into account this embedded constraint under *SPCImage*, FLI-net is ≅ 30 times faster than *SPCImage* per pixel processed (FLI-Net = 9.5e-3 ms/pixel; *SPCImage* = 0.28 ms/pixel).

**Figure 3.**
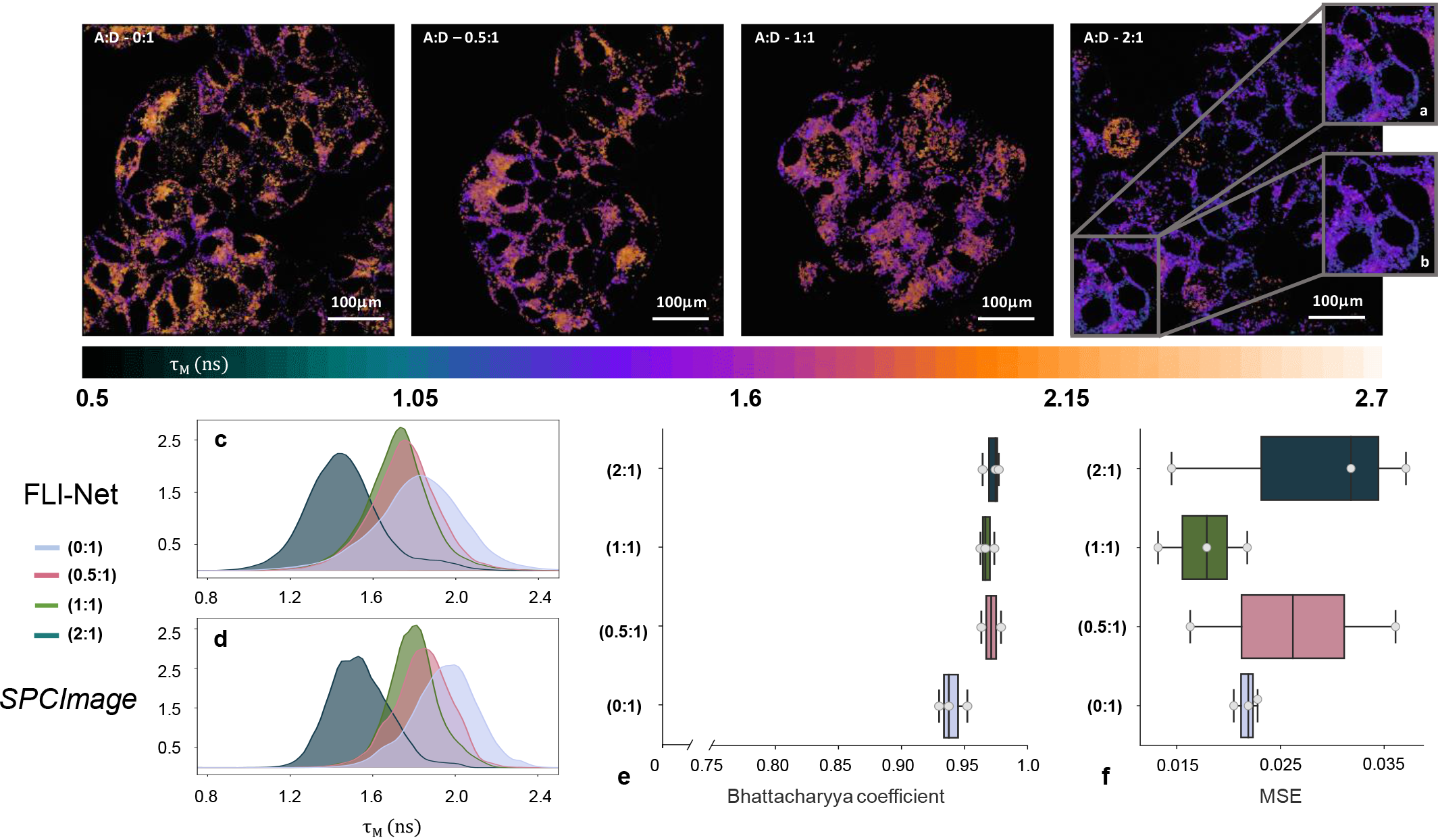
Visible FLIM microscopy data. Representative ***τ***_***m***_ maps obtained via FLI-Net using T47D cells containing Tf-AF488 (A:D = 0:1) or different donor-acceptor ratios of Tf-AF488 and Tf-AF555 (0.5:1, 1:1 and 2:1). **a-b**, Representative ROI comparison between FLI-Net (**a**) and *SPCImage* (**b**). **c-d**, Distribution histograms of ***τ***_***m***_ obtained via FLI-Net compared to *SPCImage*. **e-f**, Bhattacharyya coefficient and MSE results for three microscopy voxels at each A:D ratio.

**Figure 4.**
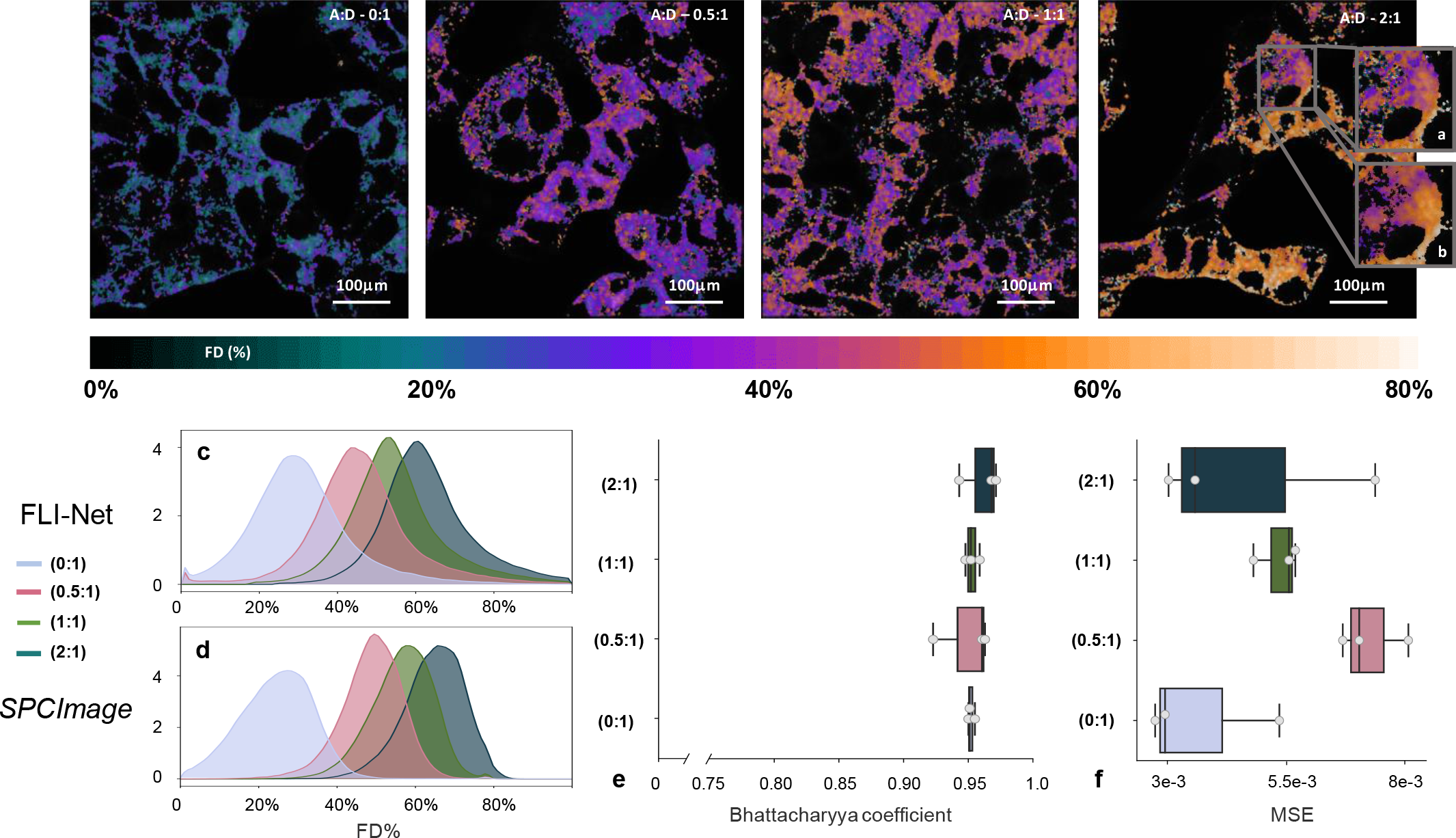
NIR TCSPC FLIM microscopy data. Representative FRET-Donor percentage maps obtained via FLI-Net using T47D cells containing Tf-AF700 (A:D = 0:1) or A:D ratios of Tf-AF700 and Tf-AF750 (0.5:1, 1:1 and 2:1). a-b, Example ROI comparison between FLI-Net (a) and *SPCImage* (b). c-d, FRET percentage distribution overlays for both techniques. e-f, Bhattacharyya coefficient and MSE results for three microscopy voxels at each volumetric fraction.

Visible FLIM is favored by relatively large lifetimes compared to the IRF temporal spread. This facilitates fitting methodologies as the IRF has minimal impact on the quantification accuracy. Though, with the impetus of translating optical molecular imaging to deep tissue imaging, great efforts have been deployed over the last two decades to develop NIR dyes. Yet, NIR dyes are typically characterized by shorter lifetimes that can be of the order of the IRF full-width at half-maximum (FWHM), rendering FLI quantification far more challenging. NIR microscopy data collected using a Zeiss LSM 880 confocal microscope equipped with same NIR FLIM detector (apparatus, fluorescence labeling, and data acquisition details described here)^37^ were used to further test FLI-Net’s robustness during in vitro NIR FLIM FRET analysis. T47D cells were incubated with Tf-AF700 and Tf-AF750 NIR FRET pairs with a range of acceptor-donor (A:D) ratio from 0:1 to 2:1. For the NIR FRET analysis and especially its *in vivo* applications, the parameter of interest is the fractional amplitude AR that reports on the fraction of donor undergoing FRET (FD%- or ***A***_***R***_)^38^. We provide in **Fig. 4**, upper panel, the estimated FD% for all cases investigated herein via FLI-Net (the corresponding *SPCImage* images are provided in the **Fig. S-D2**). As expected, as the A:D ratios increase, the FD% increases as well. Moreover, FLI-Net results, as in the previous FLIM examples, are in remarkable agreement both spatially and quantitatively with *SPCImage* results as evidenced with BCs close to ≅1 and very low MSE for all A:D ratios (see **Fig. 4(c,d)**). Similarly, to the visible FLI, FLI-net is ≅ 30 times faster than *SPCImage* per pixel processed (FLI-Net = 6.8e-3 ms/pixel; *SPCImage* = 0.21 ms/pixel). Even in challenging case of NIR dyes with short lifetimes, FLI-Net shows remarkable speed and precision in measuring FRET signal as indicative of receptor engagement in cancer cells.

## IV. MACROSCOPIC FLUORESCENCE LIFETIME IMAGING (GATED ICCD)

Another important application of FLI is in the imaging of large tissue at the macroscopic scale (MFLI). The applications hence range from high-throughput *in vitro* imaging^39^, *ex vivo*^40^ or *in vivo* tissue imaging^41^ for diagnostics, especially within the framework of optical guided surgery^42^, and preclinical studies^43^. Particularly, there is great interest in employing NIR MFLI as in this spectral window the background fluorescence is reduced, and deep tissue imaging can be performed with high sensitivity. The technology of choice to perform MFLI is gated ICCD as it provides fast acquisition speeds over a large field of view. As a tradeoff, MFLI does not provide the efficiency of TCSPC collection and is characterized by IRF of the size of the gate employed (typically 300ps or above). Hence, quantification of lifetime-based quantities can be very challenging. To demonstrate the potential of FLI-Net for MFLI based on gated ICCD (and hence its potential for widespread FLI applications), we evaluated its performance in two settings: well-plate imaging with concentration-controlled mixtures of two NIR dye mixtures and dynamic NIR-FRET *in vivo* imaging in live intact, small animals.

**Figure 5.**
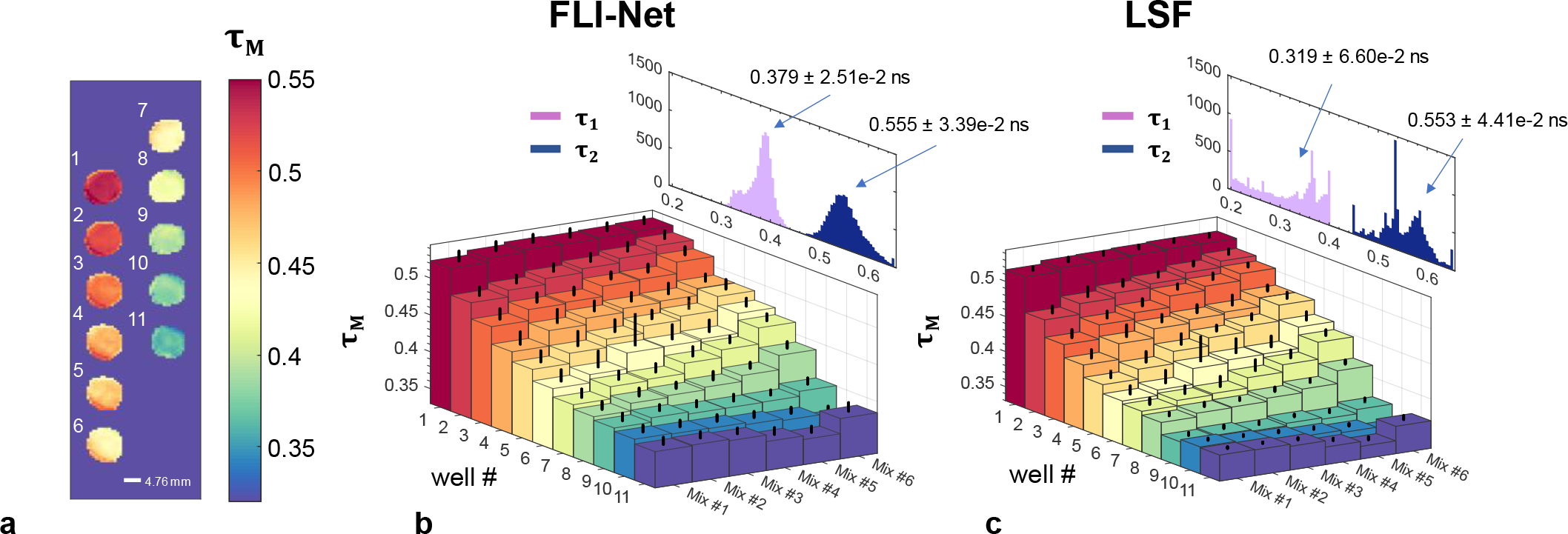
Sensitive comparison of FLI-Net with LSF via MFLI of NIR dyes possessing sub-nanosecond lifetime. **a-c**, Mean-lifetime values obtained through MFLI of six intricately prepared samples (**a**) each containing various volumetric fractions of two NIR dyes: HITCI (356 ± 4 ps) and ATTO 740 (576 ± 11 ps) in PBS buffer, both of which are excited at 740 nm and emit around 770 nm.^16^ The values obtained from the DNN (**b**) show a similar ***τ***_***m***_ trend to that of the least-squares fit (**c**) but possess a histogram that is centered much more closely to the ground-truth values of both ***τ***_**1**_ and ***τ***_**2**_.

A series of MFLI data acquired from multi-well plates, each containing a volumetric fraction of two fluorescent dyes prepared as (further described in ***Methods***), were used as a highly sensitive test of the FLI-Net’s capability to quantitatively retrieve accurate lifetimes and fractional amplitudes in controlled settings (ranging from mono-to bi-exponential). Each TPSF was captured with a time-gated, wide-field MFLI apparatus described in detail elsewhere.^44^ **Fig. 5** illustrates a sensitive comparison of FLI-Net with an LSF approach implemented in MATLAB (further described in ***Methods***). For a one-to-one comparison, the range of ***τ***_**1**_ and ***τ***_**2**_ values used for TPSF generation to train the network was set to the bounds chosen for the LSF fitting. The summary of the quantification of the two dye lifetimes (***τ***_**1**_, ***τ***_**2**_), as well as the ***τ***_***m***_ associated with the different dye ratios, for both FLI-Net and LSF are provided in **Fig. 5b/c** respectively. As can be observed, the trends exhibited for ***τ***_***m***_ are not only following the expected trendline but are also in excellent agreement between the two estimation techniques. Though, in all cases, FLI-Net provides lifetime distributions for both ***τ***_**1**_ and ***τ***_**2**_ that are centered on the expected lifetime values with a relative narrow spread.

## V. IN VIVO DYNAMICAL LIFETIME BASED IMAGING

To demonstrate the applicability of FLI-Net in a dynamic setting, we performed *in vivo* NIR FRET imaging in live and intact small animals. As demonstrated in previous studies^11,44^, the occurrence of FRET reports on the labeled Tf/TfR (ligand/receptor) engagement non-invasively. The NIR-Tf probes label the liver, as a major site of iron homeostasis regulation displaying higher levels of TfR expression. In contrast, the urinary bladder is labeled via its role as an excretion organ due to the accumulation of free dye or small labeled peptides via degradation. A total of 170 frames were acquired over a two-hour time span. Each frame consisted of 256 × 320-pixel × 160 time-gates. An experiment with a delayed injection of the acceptor compared to the donor at A:D ratio of 2:1 was performed (donor: Tf-AF700 and acceptor: Tf-AF750) as well as a FRET negative control in which only donor was injected. **Fig. 6 a/b** report on the spatially resolved FRET donor fraction (FD% or ***A***_***R***_) as estimated via FLI-Net for a few frames and for the above mentioned two conditions. In all cases the two main organs of interest, the liver and the bladder, are well resolved. Additionally, we provide the time trace of FD% over the whole 170 frames as computed for the liver and bladder ROI, both for FLI-Net and LSF (**Fig. 6c/d** and **e/f**, respectively). As expected, the FD% is reduced throughout the imaging period in the bladder as no TfR-Tf binding occurs, while the FD% increases sharply in the liver at A:D ratio of 2:1 due to abundant TfR expression in this organ. Such results are in accordance with our previous studies using the same biological system^45^. Of importance, FLI-Net provided a smoother mean FD% estimate over the organs with lower standard deviation range. Additionally, at the onset of the experiments, at which time point only the donor has been injected in the animal and hence, no FRET can occur, the baseline of FD% in the liver and bladder are similar (as expected) as estimated using FLI-Net conversely to the LSF estimates. Lastly, FLI-Net demonstrated these remarkable performances at speeds readily employable for real-time use, ≅ 32 ms/frame versus ≅ 7.5 × 10^6^ ms/frame with assistance of a binary mask for the LSF.

**Figure 6.**
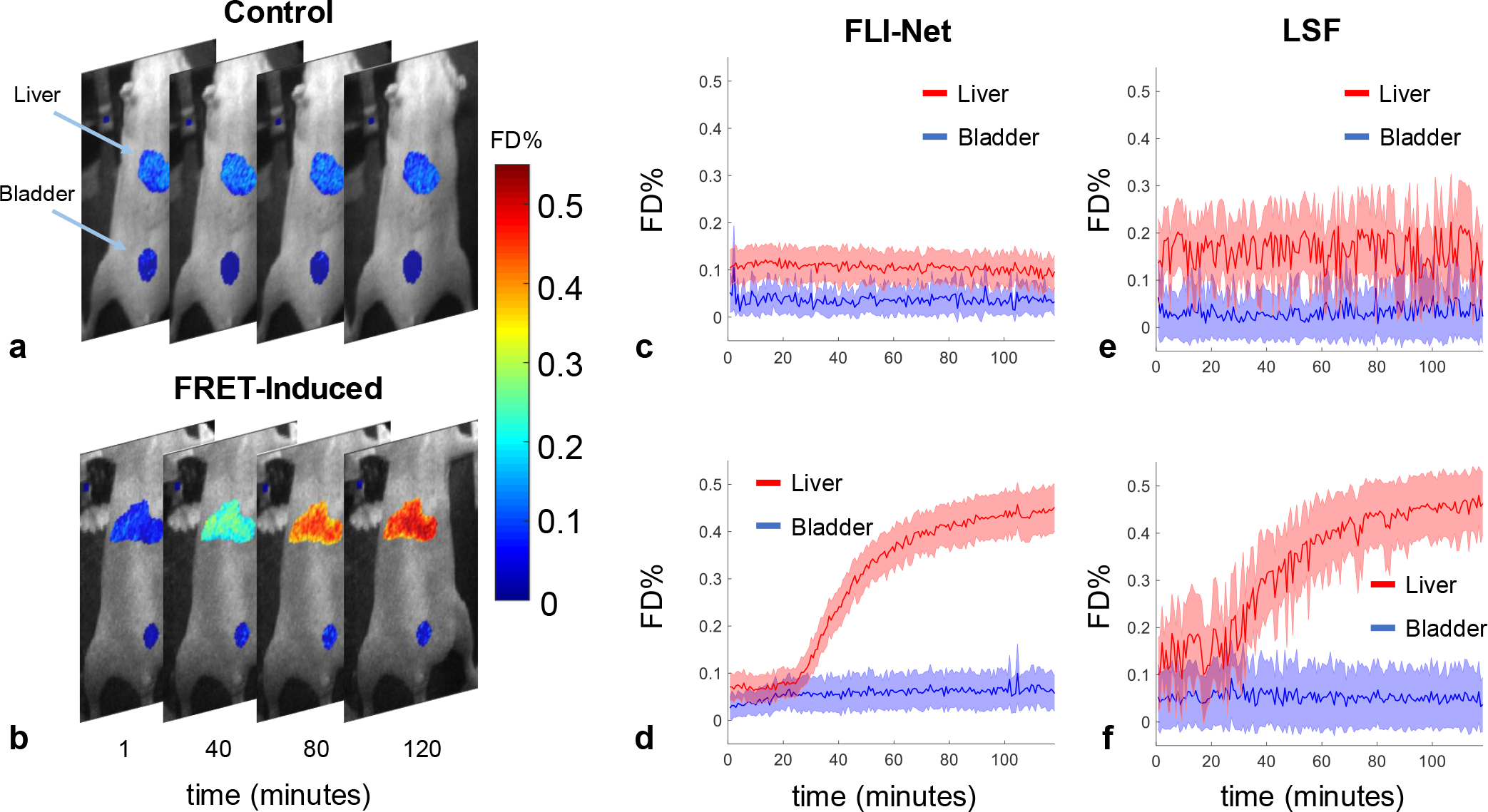
Dynamical FRET-MFLI performed over a two-hour time span. Four equally-spaced (in-time) MFLI-FRET image overlays obtained from two mice: tail-vein injected Tf-AF700 (donor-only control) (**a**) and tail-vain injected Tf-AF700 followed 20 min later by injection of Tf-AF750 (**b**). Imaging was performed over a time-span of approximately 2 hours after Tf-AF700 injection (further detailed in the *Supplementary Materials*). **c-f**, comparison of LSF versus FLI-Net results for FRET donor percentage (∝ Tf/TfR engagement). The shaded region associated with each curve corresponds to the standard deviation of all values obtained for both the liver and urinary bladder at each time-point. The computation time required for the LSF was > 2 hours using only the masked regions (liver and urinary bladder), whereas FLI-Net produced all ~ 170 parameter maps, using the entire 256 × 320-pixel acquisition, in < 48 seconds (~ 32 ms/voxel).

## VI. DISCUSSION

FLI imaging is a popular technique that enables accurate probe quantification in biological tissues revealing unique information of great value for the biomedical community. To derive the lifetime-based quantities, model-based methodologies or graphical approaches have been proposed but they rely on relatively complex inverse formulation, may necessitate calibration samples and/or need to be adapted to the application investigated and instrumentation employed. In contrast, Deep Learning methodologies can deliver ultra-fast and parameter-free processing performance. Our proposed 3D CNN architecture, as well as training methodology, offers the potential of a generalized tool for ultrafast, parameter/fit- free, quantitative FLI imaging for a wide range of applications and technologies. Especially, our methodology is centered on model-based learning that is efficient, robust and accurate. Such approach avoids the need to acquire massive training datasets experimentally and can encompass numerous applications and/or technologies.

Beyond the unique features of FLI-Net, our FLIM and MFLI studies establish our analytic framework as a robust and quantitatively accurate tool for FLI studies over a large range of lifetimes (visible-NIR), photon count and technologies employed. Of particular note, FLI-Net was robust in estimating lifetime-based parameters both in the cases of mono- and bi-exponential decays without setting any parameters (see **Table S4/S5** and **Fig. S9**) As FLI-Net was trained using both these models, there was no need to have it trained on a specific case, whereas, fit techniques require the user to select the model based on its preferences/expectations. Hence, FLI-Net is free of such bias. Additionally, we have selected to benchmark FLI-Net versus the commonly employed fitting software in the field.^47^ However, it is important to note that such methodologies are well known to perform poorly in the cases of low photon counts. This was also a fundamental limitation in a previous study proposing to use a basic Artificial Neural Network, ANN-FLIM^46^ that was not able to retrieve the life-time based parameters in all cases, especially with low p.c. Conversely, FLI-Net produced robust estimate in such conditions as depicted in **Fig. 1d** (p.c._low_∈[25,100]). These results indicate that FLI-Net performs well in photon starved conditions which are prevalent in biological studies. Last, FLI-Net is an approach that estimates, by design, the lifetime-based parameters for whole image at once. The computational time reported herein are hence for the full FLI acquisition and not limited to an ROI, as typically done for fitting techniques. Moreover, in the case of MFLI, the computational times were enabling real-time live animal imaging (≅ 32 ms/frame). Hence, FLI-Net is well positioned to profoundly impact clinical applications such as fluorescence guided surgery currently employed in a few scenarios. In this context, FLI is expected to play a major role either by providing unique contrast mechanisms such in pH-transistor like probes^42^ or improve sensitivity for current clinically approved dyes^43^. However, to date, FLI imaging formation was not attainable at speed relevant to current clinical practice, but FLI-Net can overcome this important barrier for clinical translation.

Herein, we have demonstrated the quantitative accuracy, as benchmarked against current methods, both in microscopic and preclinical settings. Of note, all experiments provided quantities in agreement with the expected biological outcomes. For instance, FLI-Net was able to clearly discriminate between non-cancerous and cancerous cells, as well as between different types of cancer cells (AU565 and T47D vs. MDA-MB-231). In contrast to *SPCImage*, FLI-Net reported on a significant difference between the ***τ***_***m***_ of untreated and Na cyanide across all cell types (**Fig. 2i**). In the case of the cell line AU565, *SPCImage* quantified an increase in metabolic status after Na cyanide exposure conversely to FLI-Net and the expected effect of this metabolic inhibitor. Moreover, a crucial advantage of FLI-Net vs. *SPCImage* is that it takes into consideration all pixels in the image instead of relying on biased ROI selection. Beyond the overall image quantification as reported, it is important to note also that FLI-Net images show punctate distribution of Tf-containing endosomes and heterogeneity of Tf uptake across cancer cells. This illustrates the potential of FLI-Net to report on important biological information without requiring any parameter settings or ROI selection by the users. Such findings in the microscopic settings are also confirmed in the preclinical studies in which the FRET Donor fraction (FD%) prior to the delayed injection of the Acceptor were in accordance with expectation for FLI-Net but overestimated when using LSF in the case of the liver. Taken altogether, these results suggest that FLI-Net provides highly reliable results with increased sensitivity and reproducibility compared to current iterative methodologies.

Beyond the topical application of fit-free FLI, the overall architecture of FLI-Net and the associated training methodology has potential for application across a myriad of biomedical imaging techniques that currently utilize a least-squares model-based fit for parameter extraction. It is common for many of these techniques to cite speed as a main hurdle they have yet to overcome for successful adoption into the clinical or commercial realm. We believe that this work provides ample supporting information and a robust proof of concept for similar adaptation and implementation in projects regarding analytic optimization across the field.

## VII. METHODS

### FLI-Net architecture and training methodology

The FLI-Net architecture consists of two main parts – 1) a shared branch focused on spatially-independent temporal feature extraction and 2) a subsequent three-junction split for independent reconstruction of ***τ***_**1**_, ***τ***_**2**_, and ***A***_***R***_ images simultaneously. Within the shared branch, spatially-independent convolutions along time (illustrated in **Fig. S1** as the blue rectangular prism with kernel size of (1 × 1×10)) was set as the network’s first layer in order maximize TPSF feature extraction. A corresponding stride of k = (1,1,5), used initially to reduce parameter count and increase computational speed, resulted in no observable decrease in performance. A residual block, possessing a kernel size of (1 × 1 × 5), followed immediately afterwards to further extract time-domain information.

To ultimately obtain image reconstruction of size (*x* × *y*) via a sequence of downsampling, a transformation from 4D to 3D was required. Thus, after the 3D-residual block (output of *x* × *y* × *n* × 50) the tensor was reshaped to dimension (*x* × *y* × (*n* × 50)), where *n* corresponds to a scalar value dependent on the number of TPSF time-points as well as the chosen network hyperparameters. This value can be determined via the following expressions:

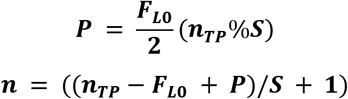

Where ***n***_***TP***_, *P*, ***F***_***L0***_, and *S* denote the number of time-points, padding (obtained through first equation), filter length along the temporal direction of the first 3D-convolutional layer (length of 10 in this study) and the corresponding stride value used in the first convolutional layer (value of 5 in this study), respectively. After this transformation, a convolutional layer of size (1×1) possessing 256 filters along with a subsequent residual block couplet possessing size (1×1) was employed before the tri-reconstruction junction. The (1×1) size of these 2D convolutional filters proved crucial in maintaining spatially-independent feature-extraction.

FLI-Net was written and trained using the machine-learning library Keras^48^ with Tensorflow^49^ backend in python. 10,000 TPSFS voxels were used during training (8000) and validation (2000), along with a batch size dependent on the target input length along time (32 for NIR, 20 for visible). Mean-squared error (MSE) was set as the loss function for each branch. The RMSprop^50^ optimizer was chosen with an initial learning rate set to 1e-6. The network was normally trained for 250 epochs using a NVIDIA TITAN Xp GPU. This training time varied slightly depending on TPSF length; ranging between 50s and 80s per epoch (for voxels possessing 160 and 256 time-points, respectively).

### Generation of the Simulation Data

For every training sample, an MNIST^51^ binary image was chosen at random and every non-zero pixel was assigned a value of intensity (*I*), short lifetime (***τ***_**1**_), long lifetime (***τ***_**2**_) and fractional amplitude (***A***_***R***_) (**Fig. S2a**). These values at each pixel, along with a randomly selected IRF (an example of which is given in **Fig. S2b** as the pink dashed line), were subsequently used in the generation of each TPSF (***Γ***(***t***)) via the equation:

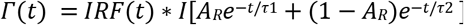

Each TPSF was normalized to a maximum intensity value of one in the last step.

### Pre-processing of gated-ICCD data

It is common for raw fluorescence time decay data to be represented initially at each time-point using a separate TIFF image. Concatenation of these along the temporal axis along with a subsequent removal of pixels possessing maximum photon counts of less than 250 was performed before the use of a Savitzky-Golay filter^52^ (length five, 3rd order) The effect of dark-noise was removed via subtraction with the mean value of time-points 1-10 (before the IRF begins ascent). Afterwards, each value was normalized to one by division with its maximum value.

Use of the Savitzky-Golay filter, or a filter that preserves the slope of the TPSF’s ascent while also having no broadening effect on the curve (as is the case with gaussian or moving average filters) was essential for analysis of the mice data given FLI-Net sensitivity along time and seemingly useful for future work regarding extraction of depth-related parameters from the TPSF. **Fig. S3** further illustrates this reasoning.

### Pre-processing of TCPSC microscopy data

Given that the TCSPC microcopy data possessed significantly lower photon counts spatially relative to MFLI, local neighborhood binning was employed. The *SPCImage* software’s pre-processing technique involves performing this binning, along with discounting any pixels possessing a maximum photon count below a specific threshold, at every pixel before fitting.^29^ This allows the software to somewhat keep the initial resolution while simultaneously fitting every pixel to a TPSF of enhanced photon count. This processing sequence was replicated for FLI-Net data sets. A maximum photon count threshold was placed initially (normally at 3 or 4), directly followed by a local neighborhood binning (5×5 kernel). Each subsequent pixel was convolved with a Savitzky-Golay filter (length 5, 3rd order). Prior work has illustrated the accurate retrieval of FRET parameters over a large variation in gate-width during MFLI as well as after introduction of artificial gating in TCSPC FLIM analysis.^53–55^ Given that the IRFs used in our data generation procedure were obtained from a gated-ICCD system, a floating average filter of length eight was introduced to mimic this effect without any reduction in temporal length. The processing methodology following steps are the same as was described in the prior section.

### NAD(P)H FLIM *in vitro*

All cell lines were obtained from ATCC (Manassas, VA, USA) and cultured in respective media at 37°C and 5% CO2. T47D and MDA-MB231 cells were grown in Dulbecco’s modified Eagle’s medium (Life Technologies) supplemented with 10% fetal bovine serum (ATCC), 4 mM l-glutamine (Life Technologies), 10 mM HEPES (Sigma). AU565 cells were cultured in RPMI medium (Life Technologies) supplemented with 10% FBS and 10 mM HEPES. MCF10A cells were cultured in DMEM/F12 medium (Life Technologies) supplemented with 5% horse serum (Life Technologies), 20 ng/mL EGF (Peprotech), 0.5 mg/mL hydrocortisone (Sigma), 10 ug/mL bovine insulin (Sigma), 100 ng/mL cholera toxin (Sigma) and 50Units/mL/50μg/mL penicillin/streptomycin (Life Technologies). For imaging experiment, the cells were plated on MatTec 35 mm glass bottom plates (Ashland, MA) 400,000 cells per plate, cultured overnight in corresponding phenol red-free complete medium and imaged in the same medium. In parallel, cells were incubated for 30 minutes with 2.5 mM NaCN in complete medium for metabolic inhibition. For FLIM imaging of NAD(P)H autofluorescence emission was performed using the Becker & Hickl HPM-100-40 detector which was attached to the NDD port on the LSM 880 using a Zeiss T-adapter that contained a 680 nm SP blocking filter (Semrock FF01-680-/SP-25 blocking edge multiphoton short-pass filter) followed by a 550/88 BP (Semrock FF01-550/88-25 single band pass filter) at spectral range of 506-594 nm. The pixel dwell time was 2.58 µs and the frame size 512 × 512 pixels. The emission was collected for 60 seconds.

### Visible FLIM-FRET *in vitro*

T47D cells were plated on MatTec 35 mm glass bottom plates as described above and cultured overnight. After that cells were washed with HBSS buffer, incubated for 30 min in DHB imaging medium (phenol red-free DMEM, 5 mg/mL bovine serum albumin (Sigma), 4 mM L-glutamine, 20 mM HEPES (Sigma) pH 7.4) to deplete native transferrin followed by 1h uptake of holo (iron-loaded) Tf-AF488 and Tf-AF555 (Life Technologies, NY) with various Acceptor: Donor ratio in DHB solution keeping Tf-AF488 concentration of 20 μg/mL constant. The uptake was terminated by washing with phosphate buffer saline and fixing in 4% paraformaldehyde. The images were acquired on Zeiss LSM 550 equipped with FLIM detector (Becker and Hickl) as described previously^56^.

### NIR FLIM-FRET *in vitro*

Human holo Tf (Sigma) was conjugated to Alexa Fluor 700 or Alexa Fluor 750 (Life Technologies) through monoreactive N-hydroxysuccinimide ester to lysine residues in the presence of 100 mM Na bicarbonate, pH 8.3, according to manufacturer’s instructions. T47D cells were processed for Tf uptake in the same manner as described above. NIR FRET FLIM was performed on Zeiss LSM 880 Airyscan NLO multiphoton confocal microscope using a HPM-100-40 high speed hybrid FLIM detector (GaAs 300-730 nm; Becker & Hickl) and a Titanium: Sapphire laser (Ti: Sa) (680-1040 nm; Chameleon Ultra II, Coherent, Inc.). The Ti: Sa laser was used in conventional one-photon excitation mode. Because of this, the FLIM detector must be attached to the confocal output of the scan head. On the LSM 880 with Airyscan, the confocal output from the scan head was used for the ‘Airy-Scan’ detector and thus it was not directly accessible. However, to accommodate the HPM-100-40 detector a Zeiss switching mirror was inserted between the scan head and the Airyscan detector. The 90° position of the switching mirror directs the beam to a vertical port to which the FLIM detector was attached via a Becker & Hickl beamsplitter assembly. A Semrock FF01-716/40 band pass filter and a FF01-715/LP blocking edge short-pass filter were inserted in the beamsplitter assembly to detect the emission from Alexa 700 and to block scattered light, respectively. The 80/20 beamsplitter in the internal beamsplitter wheel in the LSM 880 was used to direct the 690nm excitation light to the sample and to pass the emission fluorescence to the FLIM detector.

### NIR MFLI well-plate series

To test the sensitivity of FLI-Net for extracting bi-exponential parameters, we mixed two NIR dyes, ATTO740 (A740, 91394-1MG-F, Sigma-Aldrich, MO) and 1,1′,3,3,3′,3′-hexamethylindotricarbocyanine iodide (HITCI, 252034-100MG, Sigma-Aldrich, MO) initially prepared in PBS at various initial concentrations (**Table S2**). For each concentration pair, different volumes of both dyes were mixed to obtain a total volume of 300 μL with volume fractions ranging for 0 to 100% (10% steps, see **Table S3**).

### Dynamic NIR MFLI-FRET *in vivo*

In these experiments, the dynamics of FRET were observed by injecting Tf probes labeled with donor and acceptor at different time points. For all experiments, athymic nude female mice (Charles River, MA) were first anesthetized with isoflurane (EZ-SA800 System, E-Z Anesthesia), placed on the imaging stage and fixed to the stage with surgical tape (3M Micropore) to prevent motion. A warm air blower (Bair Hugger 50500, 3M Corporation) was applied to maintain body temperature. The animals were monitored for respiratory rate, pain reflex, and discomfort. The mice were imaged with the time-gated imaging system in the reflectance geometry, with adaptive greyscale illumination to ensure appropriate dynamic range between the regions of interest. In particular, excitation intensity had to be reduced in the urinary bladder due to accumulation of NIR-labeled Tf over time. Two hours after tail-injection of 20 μg of Tf-AF700, the FRET-induced mouse was imaged for ~15 minutes before retro-orbital injection with 40 μg of Tf-AF750 (A:D ratio 2:1). Imaging was continued for another 105 minutes. For the negative control mouse (0:1), no further probe was injected throughout the imaging session. **Fig. S10** is provided for further clarity. The time-resolved MFLI-FRET imaging system used in this study was described in detail elsewhere^44^.

### LSF Analysis

The LSF implementation chosen for use was based around the MATLAB’s function *fmincon()*^57^. The lower and upper bounds of both lifetime values were, for all three cases (**Fig. 1d**, **Fig. 5(e-f)**., **Fig 6f.**, **Fig. S9**) chosen to match the bounds used in generation of the TPSF data voxels utilized in training our model (**Table S1**).

### Bhattacharyya Coefficient

Given that every in vitro dataset possessed a distribution of values post-analysis, the addition of a metric for comparison of these probability distributions between FLI-Net output and *SPCImage*’s was included. To measure the degree of overlap between distributions obtained through both techniques, the Bhattacharyya coefficient was employed. Given two continuous probability distributions *M(x)* and *N(x)*, the Bhattacharyya coefficient is calculated as follows.

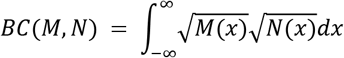

Where, when *M(x)* = *N(x)*, or, the probability distributions overlap perfectly, the Bhattacharyya coefficient is equal to 1. The metric is explained in further depth elsewhere^58^.

## Acknowledgment

We gratefully acknowledge the support of NVIDIA Corporation with the donation of the Titan Xp GPU used in this research. We would like to thank Dr. Uwe Kruger for valuable discussion and insight, Ms. Kathleen (Sez-Jade) Chen for technical support in lifetime fitting and AMC’s Imaging Core facility for the use of its confocal microscopes.

## Author contributions

J.S & X.I conceived the original idea. J.S, R.Y. & P.Y. designed the Convolutional Neural Network. J.S & X.I designed the research study. A.R, J.M & M.B acquired the FLIM data sets. N.S acquired the MFLI data sets. J.S performed the research study, the data processing and analyses of results. J.S, M.B & X.I interpreted the results. J.S & X.I wrote the manuscript. N.S., A.R., R.Y., P.Y & M.B edited the manuscript. All authors discussed the conclusions and commented on the manuscript.

