## Supplementary Material for "Ultra-fast fit-free analysis of complex fluorescence lifetime imaging via deep learning"


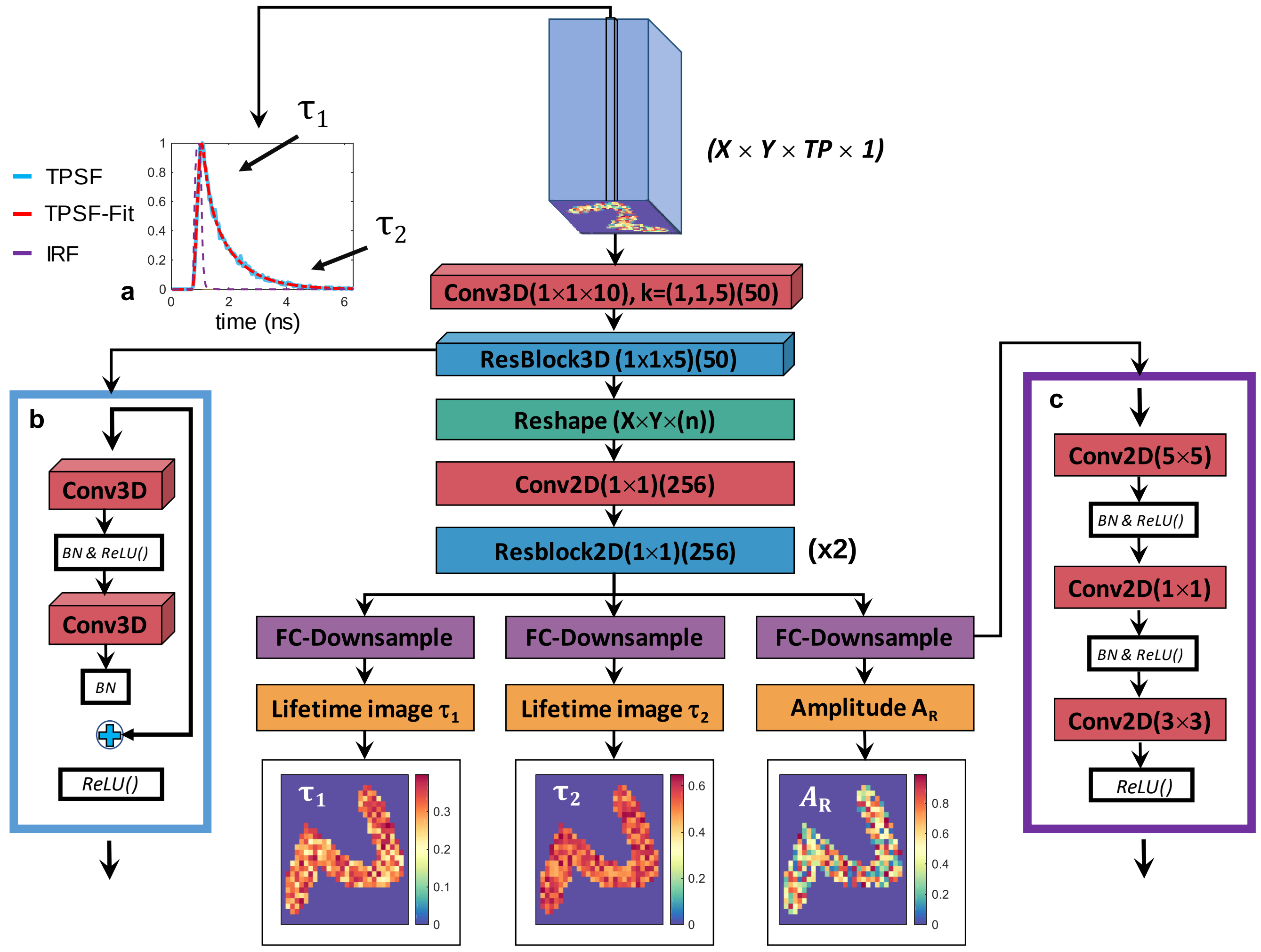


**Figure S1. Structure of "FLI-Net".** A 4D data voxel of size $X \times Y$ (pixels) $\times$ $TP$ (time-points) $\times$ 1 (**a**) is convolved with a set of 50 (1 × 1 × 10) kernels (stride represented by $k$, defaulted to one if unlisted) along the temporal axis along with a subsequent, 3D-residual block (**b**) to extract point-by-point temporal features within the earliest network layers. To obtain the three 2D maps at the end (size $X \times Y$), a reshape layer is used (depicted in green). Subsequent 2D convolutions with (1 × 1) filters downsample the parametric size without losing the fully-convolutional capability. After two 2D-convolutional residual blocks, the output is split into three independent series of fully convolutional operations for further downsampling (**c**) to simultaneously reconstruct the images of $\tau_1, \tau_2$ and $A_R$.

### Simulation Data

| Training Parameters | Fig 1 & 5 | Fig 2 & 3 | Fig 4 | Fig 6 |
|---|---|---|---|---|
| $\tau_1$ | 0.2-0.4 ns | 0.4-0.7 ns | 0.2-0.5 ns | 0.2-0.5 ns |
| $\tau_2$ | 0.45-0.65 ns | 2-3 ns | 0.9-1.1 ns | 0.9-1.1 ns |
| $A_R$ | 0-1 | 0-1 | 0-1 | 0-1 |
| **Time-Points** | 176 | 256 | 155 | 160 |
| **p.c.** | 500-2000 | 250-1500 | 250-1500 | 500-2000 |

**Table S1**. Values used for generation of TPSFs for each figure.

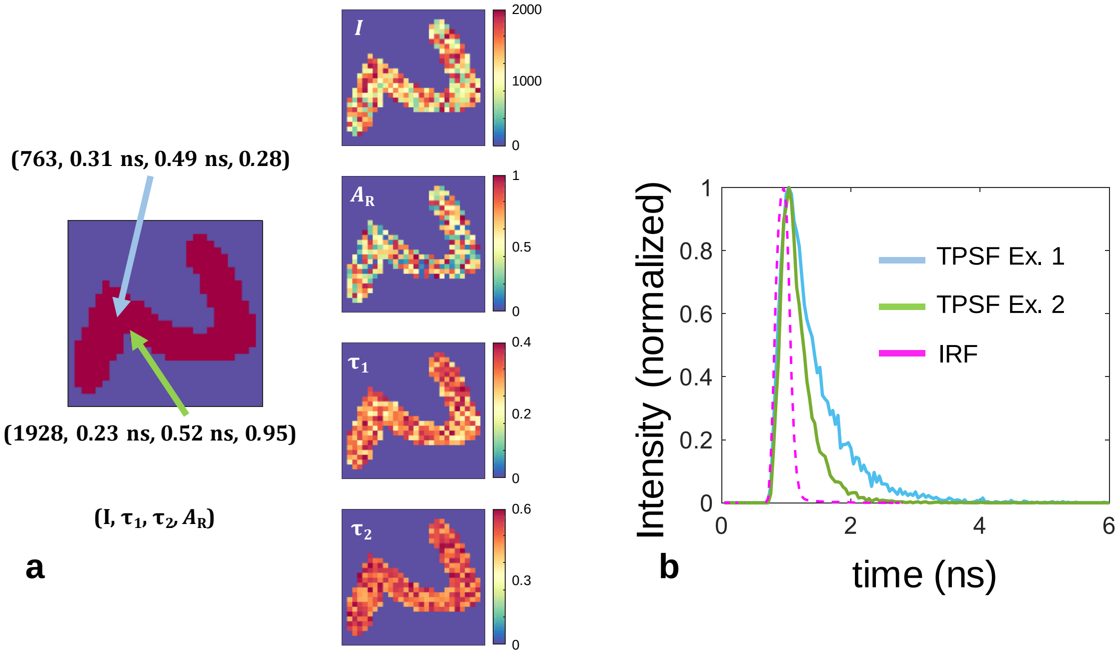


**Figure S2.** Illustration of the sequence used for creating simulation TPSF voxels used to train FLI-Net.

| Mix # | 1 | 2 | 3 | 4 | 5 | 6 |
|---|---|---|---|---|---|---|
| **Concentration ratio** | **0.5** | **0.75** | **1** | **2.5** | **7.5** | **10** |
| ATTO 740 (μM) | 10 | 10 | 10 | 10 | 10 | 10 |
| HITCI (μM) | 5 | 7.5 | 10 | 25 | 75 | 100 |

**Table S2**. Initial concentrations and concentration ratios used for preparation of each well-plate dataset used for **Fig. 5**.

| Well # | 1 | 2 | 3 | 4 | 5 | 6 | 7 | 8 | 9 | 10 | 11 |
|---|---|---|---|---|---|---|---|---|---|---|---|
| **HITCI volume fraction** | **0** | **0.1** | **0.2** | **0.3** | **0.4** | **0.5** | **0.6** | **0.7** | **0.8** | **0.9** | **1.0** |
| ATTO 740 volume (μL) | 300 | 270 | 240 | 210 | 180 | 150 | 120 | 90 | 60 | 30 | 0 |
| HITCI volume (μL) | 0 | 30 | 60 | 90 | 120 | 150 | 180 | 210 | 240 | 270 | 300 |

**Table S3**. Fluorescent dye volumes as prepared for MFLI **Fig. 5.**

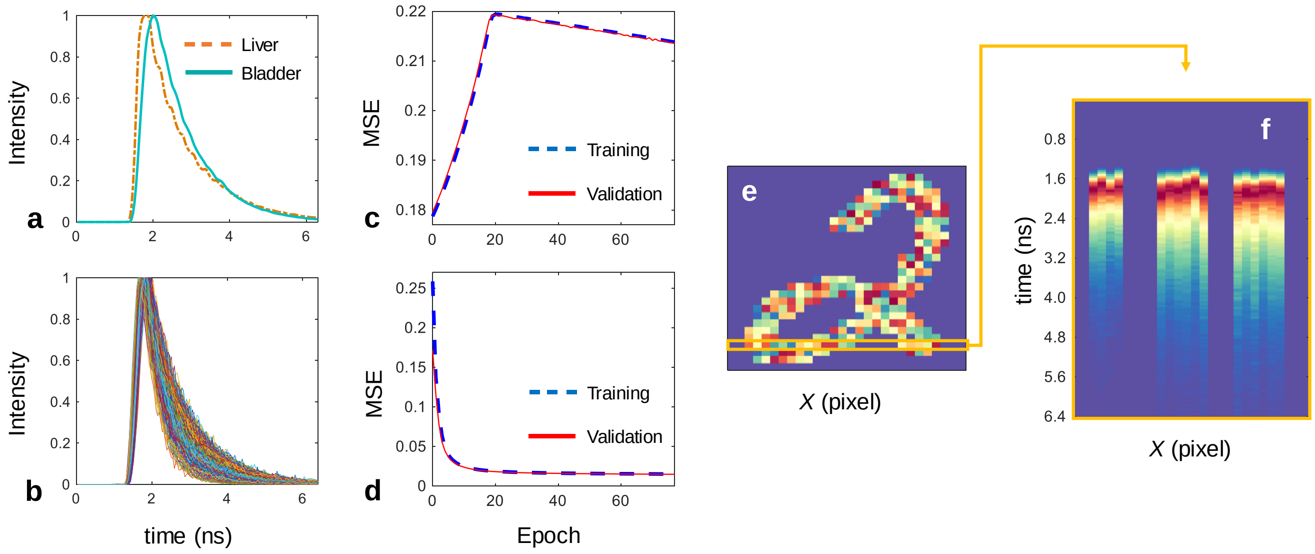


**Figure S3**. Comparison of FLI-Net with an alternative temporal-information extracting 2D-CNN architecture. **a**, Two example TPSFs obtained by averaging all TPSFs within the liver and bladder regions of a mouse during the dynamic Tf/TfR experiment at a random time-point. **b**, Example overlay of all generated TPSFs for a single simulated voxel used for training to compensate for scattering artifacts *in vivo*. The training and validation MSE curves are shown for a network architecture using only 2D, separable convolutions to extract temporal information (**c**) versus the use of 3D convolutions (**d**) along with an alternative side-view of what the generated TPSFs look like at a specific row versus time (**e-f**).

Due to various effects of scattering within different organs, *in vivo* analysis of TPSFs that do not disregard the IRF will need to account for the effects that this scattering has on the overall curve breadth. As **Fig. S3a** shows, TPSFs obtained from the bladder have a decreased slope as the curve ascends toward its maximum compared to the liver. The simulated TPSFs in **Fig. S3b** were generated by following the same procedure as previously mentioned (*Methods*: Generation of the Simulation Data) followed by convolution with a floating-average filter of random length (between 1-10) before addition of Poisson noise in order to train the DNN to be familiar with these variations in TPSF shape.

**Fig. S3c** provides the training and validation MSE curves obtained while using the data from **Fig. S3b** and a prior CNN architecture, which was developed for compressive fluorescence lifetime imaging reconstruction[6]. Separable convolutions[7] were employed with (1x1) filters for extraction of temporal information. This method of temporal feature extraction only provided high performance when the TPSFs did not possess any significant variation in ascent behavior (NIR well-plates, **Fig. 5**). **Fig. S3d** provides the training and validation MSE after switching the DNN to 3D convolutions along the temporal axis, which allowed for the results shown in **Fig. 6** and all other results given in this report.

The MSE curve behavior, illustrated in **Fig. S3c**, provides evidence that when using separable convolutions, it takes time for the network to adjust to the displacement of the TPSF's maximum value along with the various changes in slope leading up to it. The network does at around epoch 20 begin to decrease its MSE, but very slowly (plateauing after epoch 200). **Fig. S3f** provides a visualization as to how challenging it may be for a network to learn these subtle variations necessary for accurate bi-exponential fitting when faced with optical artifacts inherently inescapable in preclinical FLI application. In contrast, the use of 3D convolutions across a 4D TPSF input voxel are unaffected by the displacement of the curve maximum or the breadth of the curve, which is supported by the MSE curves in **Fig. S3d**. This is promising, especially since it would be simple to implement a network with a fourth parameter corresponding to the scattering properties, or, the randomly chosen length of the moving average filter used at that pixel location. It strongly suggests that the same network architecture would be able to learn added parameters exploiting this effect, which will be explored in future works.

### FURTHER COMPARATIVE METRICS/MATERIAL

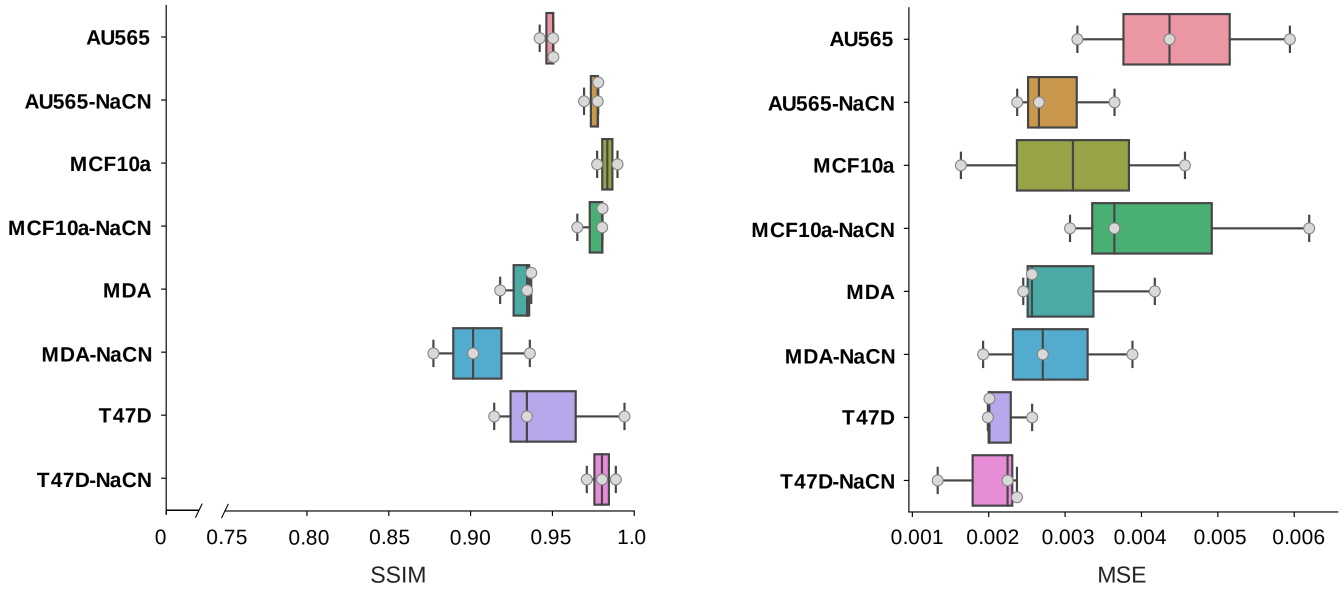


**Figure S4.** SSIM and MSE comparison results between FLI-Net and *SPCImage* $\tau_m$ images of NAD.

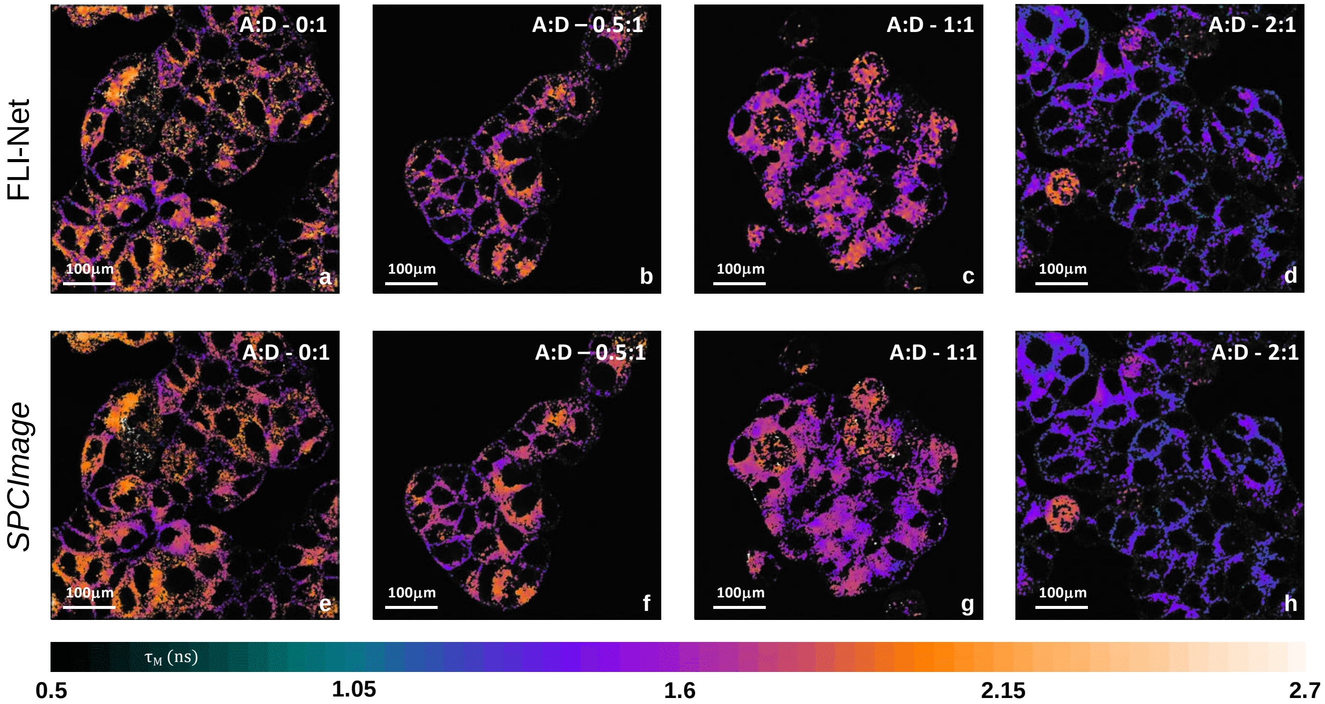


**Figure S5.** Full-image comparisons of FLI-Net vs. *SPCImage* using visible TCSPC FLIM microscopy data.

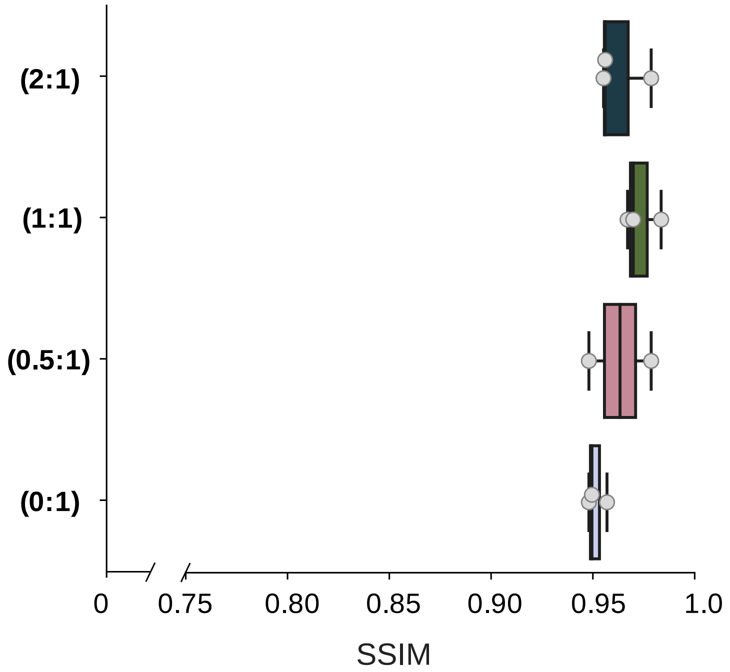


**Figure S6.** SSIM comparisons of FLI-Net vs. *SPCImage* using visible TCSPC FLIM microscopy data.

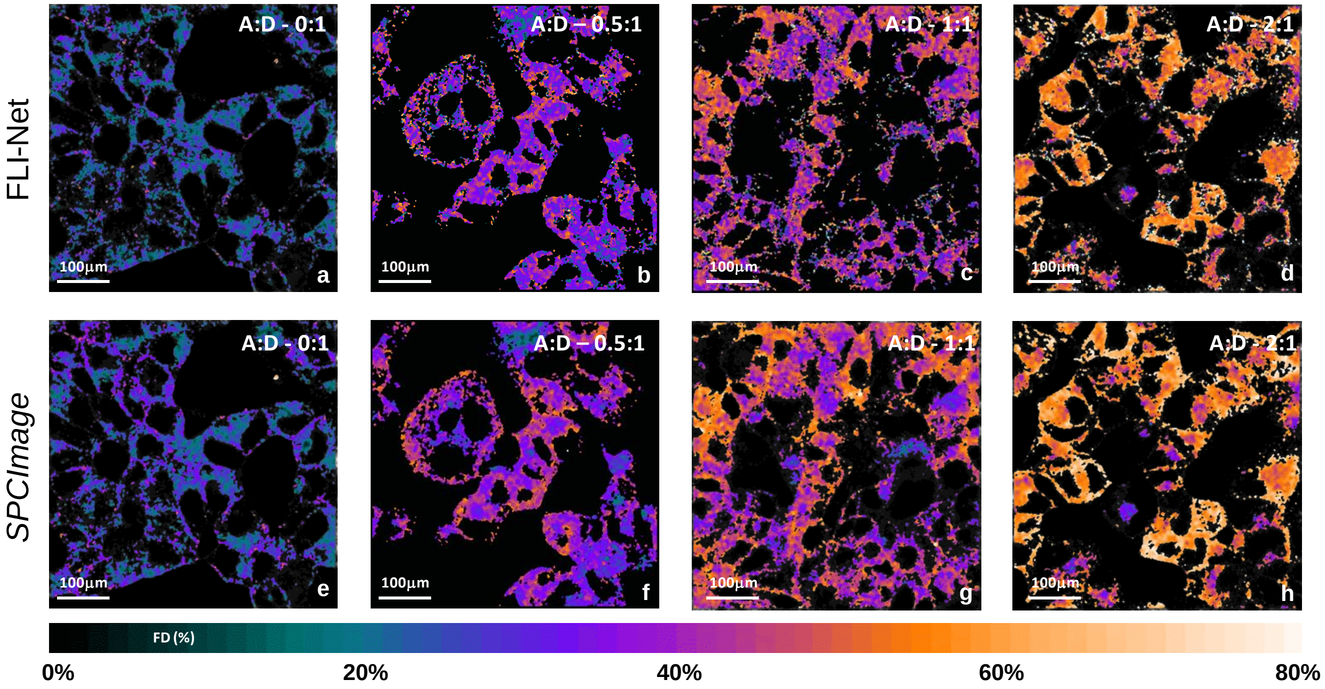


**Figure S7.** Full-image comparisons of FLI-Net vs. *SPCImage* using NIR TCSPC FLIM microscopy data.

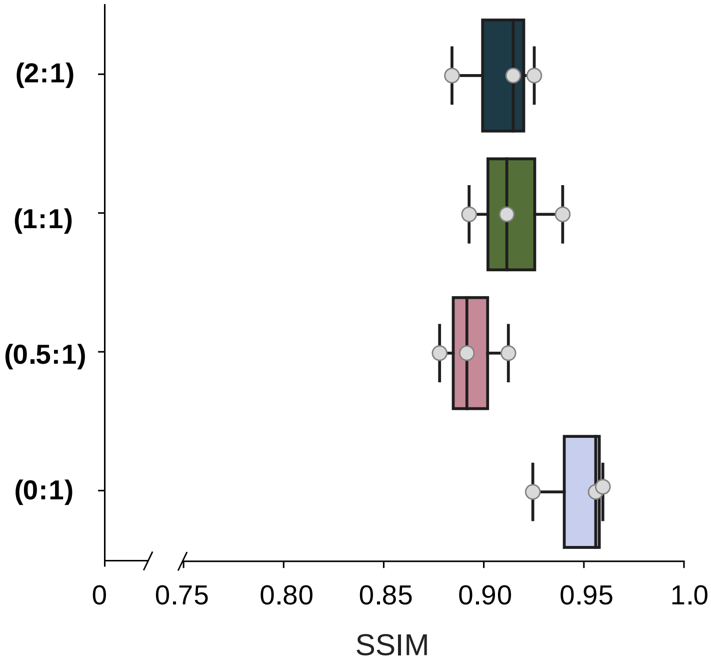


**Figure S8.** SSIM comparisons of FLI-Net vs. *SPCImage* using NIR TCSPC FLIM microscopy data.

### Mono-exponential Comparison

**Simulated data.** To assess the 3D-CNN's capability in accurately identifying mono-exponential decays, two tests were performed. Two different sets of TPSF data were simulated between differing bi-exponential lifetime and intensity bounds. The absolute errors were calculated using only pixels possessing ground-truth $A_R$ values of either below 0.05 (bottom 5%) or above 0.95 (top 5%) for both FLI-Net and LSF.

| Threshold | **FLI-Net** | **LSF** |
|---|---|---|
| $A_R$ < 5% (*n* = 7460) | 0.0437 ± 0.0372 | 0.259 ± 0.112 |
| $A_R$ > 95% (*n* = 7415) | 0.0135 ± 0.0124 | 0.0684 ± 0.0529 |

**Table S4. Average absolute error |Predicted – G.T.| obtained using both techniques**. *n* corresponds to number of total pixels evaluated (500 simulated TPSF voxels, $(\tau_1, \tau_2, p.c.)$ = ([0.2-0.4] ns, [0.45-0.65] ns, [250-500])).

| Threshold | **FLI-Net** | **LSF** |
|---|---|---|
| $A_R$ < 5% (*n* = 6323) | 0.0353 ± 0.0294 | 0.0480 ± 0.0545 |
| $A_R$ > 95% (*n* = 6393) | 0.0136 ± 0.0136 | 0.0684 ± 0.0529 |

**Table S5. Average absolute error |Predicted – G.T.| obtained using both techniques**. *n* corresponds to number of total pixels evaluated (450 simulated TPSF voxels, $(\tau_1, \tau_2, p.c.)$ = ([0.35-0.6] ns, [0.9-1.1] ns, [500-2000])).

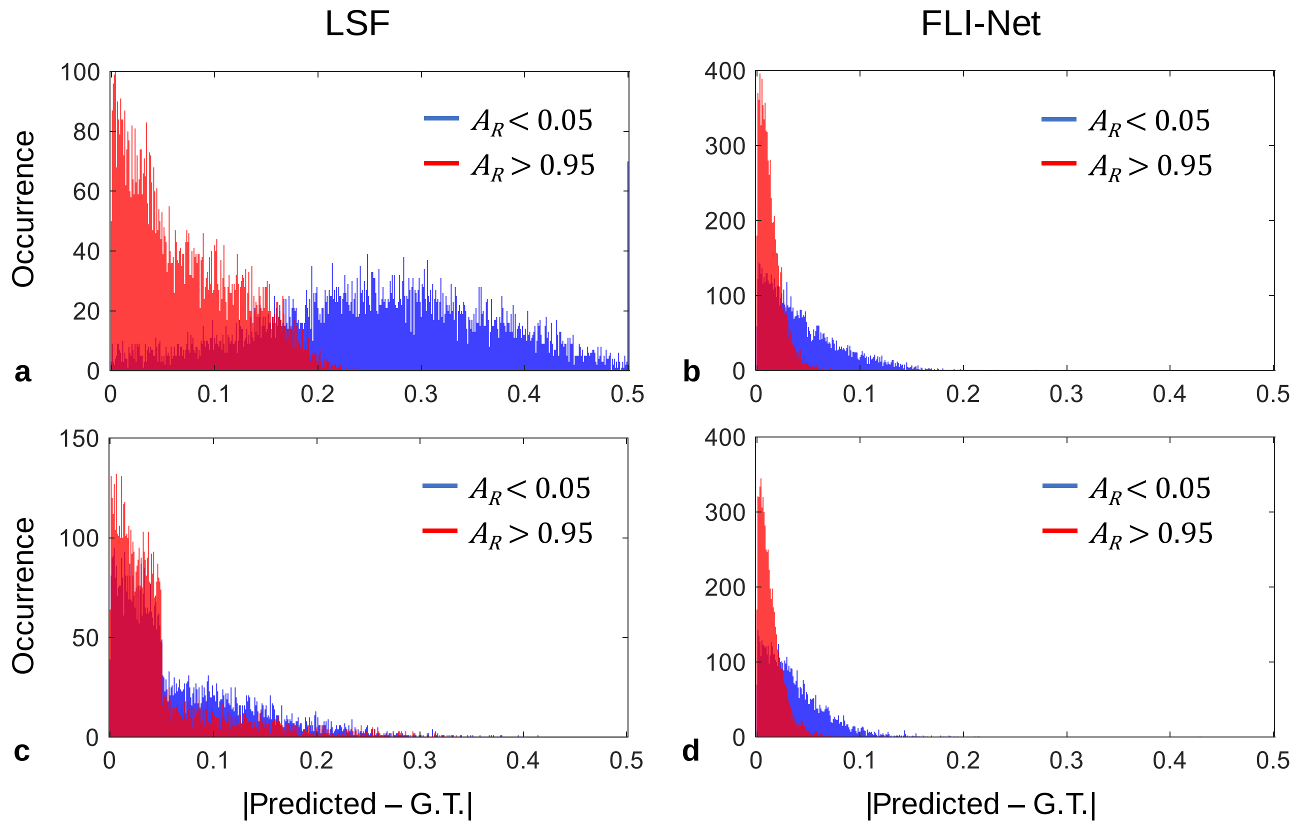


**Figure S9.** Histograms of $A_R$ absolute error. **a-b**, Absolute error for both techniques using $(\tau_1, \tau_2, p.c.)$ = ([0.2-0.4] ns, [0.45-0.65] ns, [250-500])). **c-d**, Absolute error for both techniques using $(\tau_1, \tau_2, p.c.)$ = ([0.35-0.6] ns, [0.9-1.1] ns, [500-2000])).

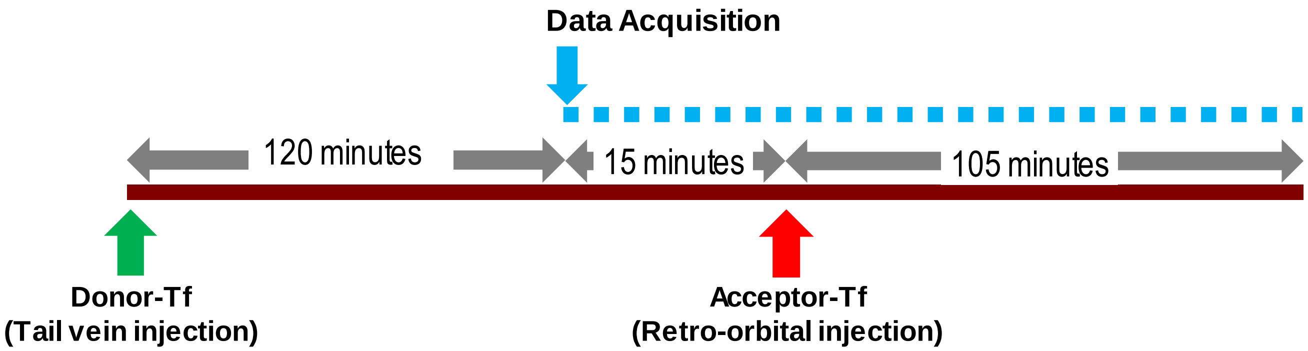


| Mouse | Donor-Tf | Acceptor-Tf |
|---|---|---|
| Control | 20 µg | - |
| FRET | 20 µg | 40 µg |

**Figure S10.** Diagram illustrating the timeline of *in vivo* MFLI acquisition and injection (Fig. 6).
